# The temperature-dependent conformational ensemble of SARS-CoV-2 main protease (M^pro^)

**DOI:** 10.1101/2021.05.03.437411

**Authors:** Ali Ebrahim, Blake T. Riley, Desigan Kumaran, Babak Andi, Martin R. Fuchs, Sean McSweeney, Daniel A. Keedy

## Abstract

The COVID-19 pandemic, instigated by the SARS-CoV-2 coronavirus, continues to plague the globe. The SARS-CoV-2 main protease, or M^pro^, is a promising target for development of novel antiviral therapeutics. Previous X-ray crystal structures of M^pro^ were obtained at cryogenic temperature or room temperature only. Here we report a series of high-resolution crystal structures of unliganded M^pro^ across multiple temperatures from cryogenic to physiological, and another at high humidity. We interrogate these datasets with parsimonious multiconformer models, multi-copy ensemble models, and isomorphous difference density maps. Our analysis reveals a temperature-dependent conformational landscape for M^pro^, including mobile solvent interleaved between the catalytic dyad, mercurial conformational heterogeneity in a key substrate-binding loop, and a far-reaching intramolecular network bridging the active site and dimer interface. Our results may inspire new strategies for antiviral drug development to counter-punch COVID-19 and combat future coronavirus pandemics.

**Synopsis:** X-ray crystallography at variable temperature for SARS-CoV-2 M^pro^ reveals a complex conformational landscape, including mobile solvent at the catalytic dyad, mercurial conformational heterogeneity in a key substrate-binding loop, and an intramolecular network bridging the active site and dimer interface.

## Introduction

COVID-19 is a global pandemic disease caused by severe acute respiratory syndrome coronavirus 2. SARS-CoV-2 is a highly infectious, airborne, respiratory virus, which has caused over 200 million infections and nearly 5 million deaths worldwide as of October 2021. Over the the past year and a half, several approaches to prevent and treat COVID-19 have been successfully developed, including new vaccines, monoclonal antibody treatments (Baum *et al*., 2020), and repurposed existing therapeutics (Beigel *et al*., 2020; Boras *et al*., 2021). However, development of novel small-molecule antiviral drugs has lagged behind. New antiviral drugs would not only provide a powerful weapon against COVID-19 for infected patients and frontline workers, but would also aid in preparation for future coronavirus pandemics.

A promising target for potential new antiviral drugs against SARS-CoV-2 is a chymotrypsin-like protease known by several names: non-structural protein 5, nsp5, 3C-like protease, 3CL^pro^, main protease, or M^pro^. M^pro^ is part of a polyprotein encoded by the viral RNA genome. After being excised from the polyprotein by its own proteolytic activity, M^pro^ cleaves at no fewer than 11 sites in the polyprotein to generate individual functional proteins (V’kovski *et al*., 2021) that help the virus replicate. Due to its importance to the SARS-CoV-2 life cycle, M^pro^ has been identified as a key target for COVID-19 drug design.

Drug design efforts focused on M^pro^ have been aided by insights from structural biology. The first SARS-CoV-2 M^pro^ crystal structures were released in the Protein Data Bank (PDB) (Berman *et al*., 2000) early in the pandemic, within the first week of February 2020 (Jin *et al*., 2020). These structures revealed that, like SARS-CoV M^pro^ before it, SARS-CoV-2 M^pro^ is composed of two β-barrel domains known as domain I and domain II, and an α-helical bundle known as domain III (**Fig. 1a**). The active site cavity is located on the surface, with the His41–Cys145 catalytic dyad positioned between domain I and domain II. Domain III is involved in regulating dimerization (Zhang *et al*., 2020), which is critical for coronavirus M^pro^ catalytic activity (Fan *et al*., 2004; Goyal & Goyal, 2020). Since the initial structures of SARS-CoV-2 M^pro^, X-ray crystallography has been used to identify promising ligand binding sites and alternate structural states of the protein, resulting in a total of over 250 available structures. These efforts included co-crystallography with an eye toward drug repurposing (Vuong *et al*., 2020; Günther *et al*., 2021), as well as crystallographic screens of non-covalent and covalent small-molecule fragments to establish new toe-holds for *ab initio* drug design (Douangamath *et al*., 2020) which were then leveraged via a crowd-sourced process to design novel inhibitors (Chodera *et al*., 2020).

**Figure 1:**
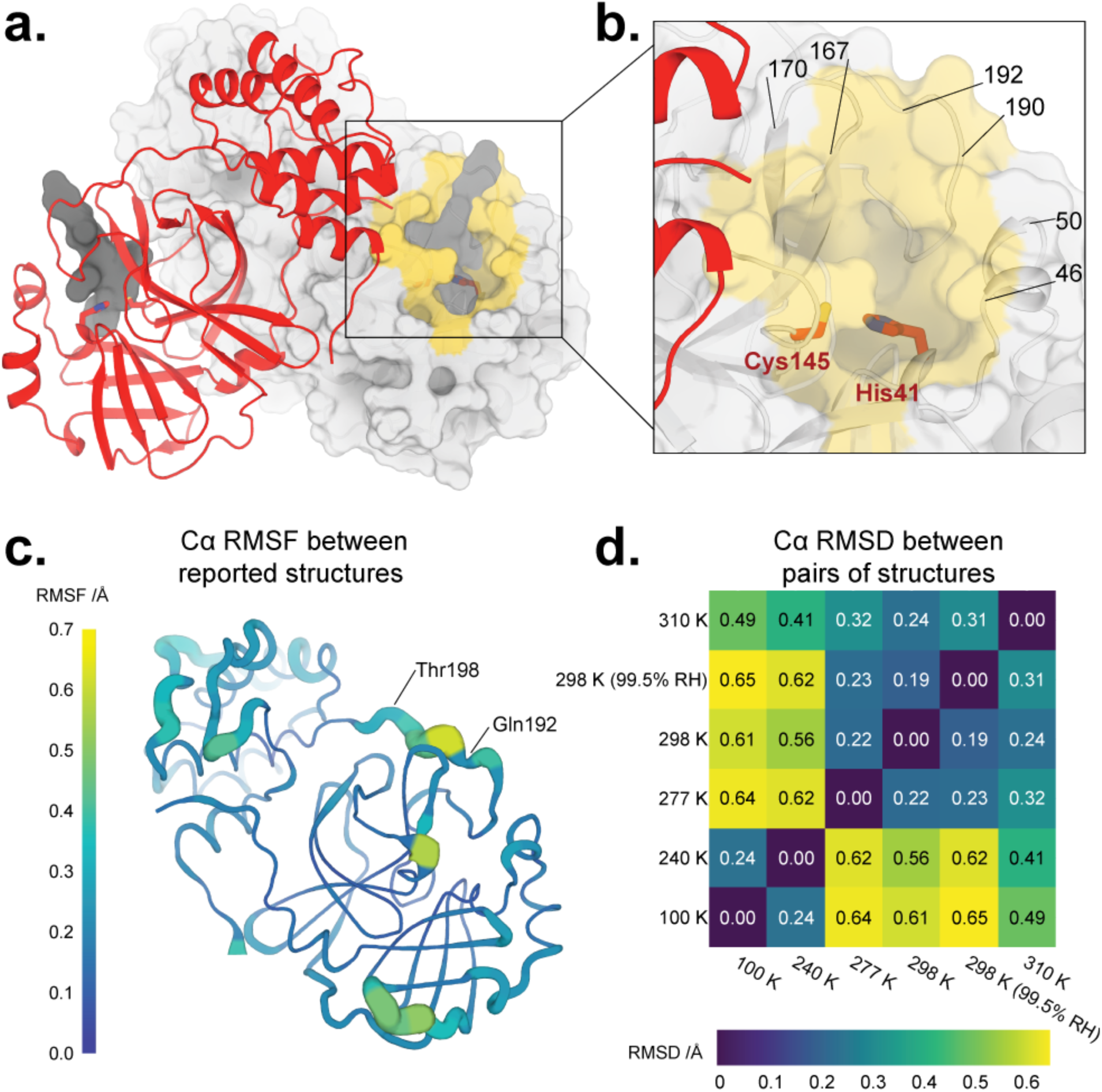
Overall structure of SARS-CoV-2 main protease at multiple temperatures. **a**. New X-ray crystal structure of apo M^pro^ at physiological temperature (310 K) (red). The biological dimer involving the other monomer (light grey surface) is constituted via crystal symmetry. The competitive inhibitor N3 from a previous structure (PDB ID 6LU7) (semi-transparent, dark grey surface) is shown in both protomers for context. **b**. Close-up view of the M^pro^ active site region, including the catalytic dyad of Cys145 and His41 (red sticks) and highlighting residues that form the substrate binding pocket (yellow surface). **c**. Cartoon putty representation of conformational variability between new M^pro^ structures described in this work: 100 K, 240 K, 277 K, 298 K, 298 K (99.5% RH), and 310 K. Thickness and color indicate root-mean-square fluctuations (RMSF) of Cα atom positions, from low (thin, dark blue) to high (thick, yellow). The largest differences between these structures’ backbones occur between residues 192–198. Same view as **a**. See also **Supp. Fig. 3**. **d**. Heatmap of pairwise Cα atom root-mean-square deviation (RMSD) between final refined structures, revealing temperature-dependent clustering (top-right vs. bottom-left). See also **Supp. Fig. 2**.

As with much modern protein crystallography, the above experiments were performed at cryogenic temperatures, which can bias protein conformational ensembles (Fraser *et al*., 2011; Keedy *et al*., 2014). To bypass this limitation, a room-temperature crystal structure of unliganded M^pro^ was reported (Kneller, Phillips, O’Neill, Jedrzejczak *et al*., 2020) (PDB ID 6WQF). Although its resolution was only moderate (2.3 Å), it nevertheless revealed evidence of conformational plasticity near the active site that was distinct relative to past cryogenic structures. Subsequent work built on this foundation of room-temperature crystallography to dissect M^pro^ function (Kneller, Phillips, Kovalevsky *et al*., 2020; Kneller, Phillips, O’Neill, Tan *et al*., 2020). However, no studies to date have reported crystal structures of M^pro^ across a wide range of temperatures. Previously, such a multitemperature crystallography strategy was instrumental for revealing novel aspects of correlated active-site conformational heterogeneity in a dynamic proline isomerase (Keedy *et al*., 2015) and of long-range allosteric signaling in a therapeutic-target tyrosine phosphatase (Keedy *et al*., 2018).

Here we report high-resolution crystal structures of SARS-CoV-2 M^pro^ at five temperatures: 100 K (cryogenic), 240 K (above the so-called glass transition or dynamical transition (Keedy *et al*., 2015)), 277 K (“room temperature” in many crystallography studies), 298 K (ambient), and 310 K (physiological). We also report a structure at ambient temperature but high relative humidity (99.5% RH) to gauge the relative effects of temperature vs. humidity on M^pro^. To our knowledge, this study represents the first experimentally based structural analysis for any SARS-CoV-2 protein at variable temperature and/or humidity. We used careful data collection with a helical strategy to minimize radiation damage, thereby isolating the effects of temperature and humidity on M^pro^. For all datasets we have constructed parsimonious multiconformer models as well as multi-copy crystallographic ensemble models, which provide complementary insights into protein structural flexibility as a function of temperature and humidity. Together, our data reveal a network of subtle but provocative temperature-dependent conformational heterogeneity, not only at the catalytic site but also spanning several functionally relevant sites throughout M^pro^, which may help motivate an allosteric strategy for antiviral drug design to combat COVID-19 and/or future coronavirus pandemics.

## Results

### Multitemperature crystallographic data collection and modeling

Data were obtained from single M^pro^ crystals using helical data collection, to maximize diffraction intensity while minimizing radiation damage (**Supp. Fig. 1**). To probe the conformational landscape of M^pro^, we obtained high-resolution structures at five different temperatures: 100 K, 240 K, 277 K, 298 K (ambient; see Methods), and 310 K. Our datasets thus span a broad temperature range: cryogenic, just above the glass transition or dynamic transition (Keedy *et al*., 2015), the range often noted as room temperature (roughly 293–300 K), and approximately physiological temperature. We also collected another 298 K dataset with high relative humidity (99.5% RH).

For all but the 277 K dataset (2.19 Å), the resolution was 2 Å or better (**Table 1**). The highest resolution was for the 100 K dataset (1.55 Å). Even at the higher temperatures, we saw little to no evidence of radiation damage (**Supp. Fig. 1**). After data reduction, we created a multiconformer model for each temperature, which includes a single conformer for most portions of the structure but alternate conformations where appropriate (Riley *et al*., 2021). See Methods for more details on data collection and modeling, and **Table 1** for overall diffraction data and refinement statistics.

**Table 1:** Crystallographic statistics for multitemperature datasets and multiconformer models. Overall statistics given first (statistics for highest-resolution bin in parentheses). RH = relative humidity. RMS = root-mean-square deviation from ideal values. For Phenix ensemble model refinement statistics, see **Table 2**.

| Structure | 100 K | 240 K | 277 K | 298 K | 298 K,<br>99.5% RH | 310 K |
| --- | --- | --- | --- | --- | --- | --- |
| PDB ID | 7MHF | 7MHG | 7MHH | 7MHI | 7MHJ | 7MHK |
| Resolution (Å) | 48.07–1.55 | 55.62–1.53 | 48.96–2.19 | 56.29–1.88 | 56.30–2.00 | 43.97–1.96 |
| Completeness (%) | 99.7 (96.0) | 100 (99.4) | 99.9 (98.7) | 100 (100) | 99.0 (97.4) | 99.9 (100) |
| Multiplicity | 3.4 (3.4) | 6.6 (6.2) | 6.9 (6.9) | 6.8 (6.9) | 6.8 (6.7) | 6.6 (6.7) |
| I/sigma(I) | 3.3 (1.0) | 7.9 (0.9) | 4.5 (1.1) | 5.0 (0.4) | 6.3 (0.6) | 5.4 (0.3) |
| R<sub>merge</sub>(I) | 0.158<br>(0.507) | 0.180<br>(1.463) | 0.292<br>(1.805) | 0.182<br>(2.353) | 0.178<br>(1.708) | 0.195<br>(1.805) |
| R<sub>meas</sub>(I) | 0.188<br>(0.604) | 1.960<br>(1.600) | 0.316<br>(1.954) | 0.197<br>(2.548) | 0.193<br>(1.854) | 0.213<br>(1.957) |
| R<sub>pim</sub>(I) | 0.100<br>(0.325) | 0.076<br>(0.639) | 0.119<br>(0.742) | 0.076<br>(0.967) | 0.074<br>(0.711) | 0.084<br>(1.046) |
| CC<sub>1/2</sub> | 0.977<br>(0.695) | 0.995<br>(0.356) | 0.985<br>(0.799) | 0.990<br>(0.285) | 0.989<br>(0.376) | 0.990<br>(0.352) |
| Wilson B-factor | 16.164 | 16.370 | 31.769 | 29.670 | 34.350 | 33.810 |
| Total observations | 127548 | 263470 | 97820 | 152368 | 125878 | 128140 |
| Unique observations | 37901 | 39975 | 14120 | 22459 | 18588 | 19444 |
| Space group | C121 | C121 | C121 | C121 | C121 | C121 |
| Unit cell dimensions (Å,Å,Å,°,°,°) | 113.71, 53.32, 44.57, 90, 102.96, 90 | 114.19, 53.49, 45.00, 90, 103.04, 90 | 115.02, 54.36, 44.97, 90, 101.50, 90 | 114.74, 54.57, 45.11, 90, 101.65, 90 | 114.88, 54.74, 45.24, 90, 101.42, 90 | 114.3, 54.29, 44.97, 90, 102.12, 90 |
| Solvent content (%) | 35.88 | 36.40 | 38.95 | 39.53 | 39.89 | 38.36 |
| R<sub>work</sub> | 0.1834 | 0.1701 | 0.1994 | 0.1847 | 0.1942 | 0.2039 |
| R<sub>free</sub> | 0.2223 | 0.2043 | 0.2547 | 0.2276 | 0.2421 | 0.2482 |
| RMS bonds (Å) | 0.006 | 0.011 | 0.002 | 0.003 | 0.004 | 0.003 |
| RMS angles (°) | 0.888 | 1.094 | 0.472 | 0.569 | 0.702 | 0.576 |
| Ramachandran outliers (%) | 0.33 | 0.33 | 0.33 | 0.00 | 0.33 | 0.66 |
| Ramachandran favoured (%) | 97.70 | 98.36 | 96.71 | 97.37 | 96.38 | 97.37 |
| Clashscore | 3.81 | 2.42 | 2.31 | 1.89 | 2.31 | 2.73 |
| MolProbity score | 1.24 | 1.02 | 1.22 | 1.07 | 1.25 | 1.18 |

### Overall structure as a function of temperature

The global structure of M^pro^ in our crystals remains similar across temperatures (**Fig. 1d, Supp. Fig. 2**), as expected. Indeed, the maximum Cα RMSD between any pair of structures in the ambient-humidity multitemperature series is only 0.64 Å, and the maximum all-atom RMSD is only 0.98 Å. However, there is a clear clustering between lower-temperature (240, 277 K) and higher-temperature (277, 298, 310 K) structures, based on either Cα RMSD (**Fig. 1d**) or all-atom RMSD (**Supp. Fig. 2**). These observations indicate that aspects of the M^pro^ conformational landscape change in response to temperature.

Humidity also appears to have some effect on M^pro^ structure, as evidenced by the fact that the overall largest pairwise Cα RMSD (0.65 Å, **Fig. 1d**) and all-atom RMSD (1.04 Å, **Supp. Fig. 2**) involve the 298 K high-humidity (99.5% RH) structure. However, the corresponding RMSD values for the 298 K ambient-humidity (36.7% RH) structure are only slightly smaller (<0.1 Å difference). These RMSD differences between high vs. low humidity are minor compared to the differences between the high vs. low temperature clusters mentioned above. Thus, temperature affects M^pro^ structure noticeably more than does humidity. This result contrasts with previous studies of lysozyme in which similar protein structural alterations were achieved by either small changes in humidity or large changes in temperature (Atakisi *et al*., 2018); this discrepancy may result from different protein:solvent arrangements in the lysozyme vs. M^pro^ crystal lattices.

### Temperature dependence of local alternate conformations

To provide more detailed insights into the observed global temperature dependence, we sought to identify alternate conformations at the local scale that were stabilized or modulated by the temperature shifts in our experiments. We specifically focused our attention on areas of the protein that are of interest for drug design and/or biological function: the active site, nearby loops associated with substrate binding, and the dimer interface.

First, the M^pro^ active site structure remains mostly consistent across our temperature series (**Fig. 2**). The catalytic amino acids are in very similar conformations across temperatures. Additionally, a key active-site water molecule (known as H_2_O_cat_), which hydrogen-bonds to both of the side chains of the catalytic dyad (His41 and Asp187), remains in the same position across our structures (**Fig. 2**). It has been suggested (Kneller, Phillips, O’Neill, Jedrzejczak *et al*., 2020) that this water may play the role of a third catalytic residue (in addition to the catalytic dyad of His41 and Cys145). As previously noted (Kneller, Phillips, O’Neill, Jedrzejczak *et al*., 2020), H_2_O_cat_ is not modeled in some cryogenic structures but it is modeled in 89% (224/252) of the publicly available structures of SARS-CoV-2 M^pro^ as of Oct. 11, 2021 (the vast majority of which are cryogenic), and perhaps should have been modeled in others (Jaskolski *et al*., 2021).

**Figure 2:**
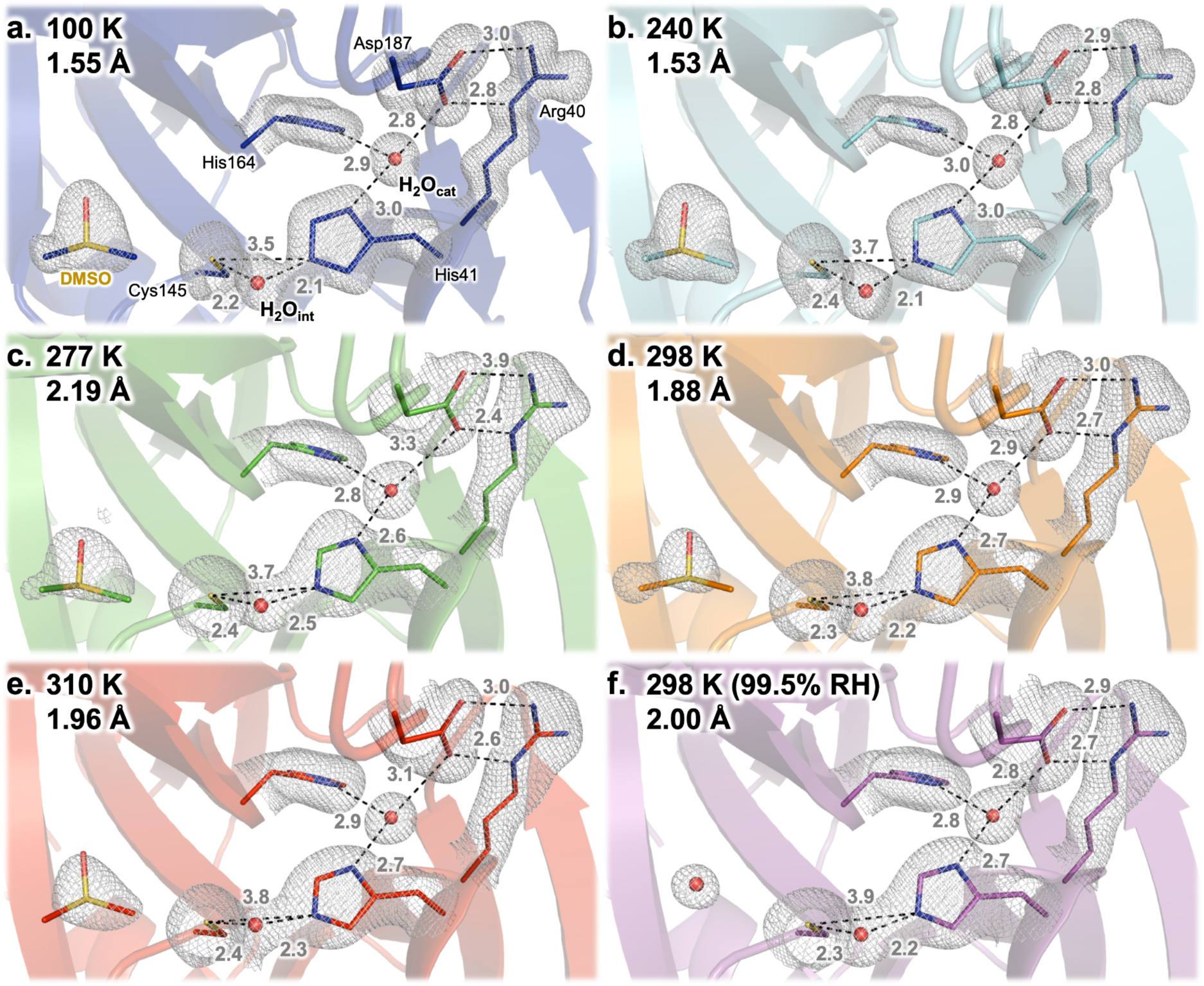
The unliganded M^pro^ active site as a function of temperature and humidity. 2F_o_-F_c_ electron density (1.0 σ, gray mesh) and interatomic distances (pink, in Å) shows that the active-site structure remains similar across datasets, including the catalytic dyad of His41 and Cys145 and the presumed catalytic water (H_2_O_cat_). One minor exception is a different water, H_2_O_int_, which tends to shift upward in this view as temperature increases, adjusting its interactions with His41 and Cys145 (see also **Fig. 3**). An ordered DMSO molecule from the crystallization solution is visible at the left of each panel, except for 298 K at high humidity (99.5% RH) in which case a water is present at the same site instead.

As with the active-site amino acids and H_2_O_cat_, a dimethyl sulfoxide (DMSO) molecule from the crystallization solution is ordered nearby in each structure in the multitemperature series (**Fig. 2**, left of each panel). Interestingly, however, this DMSO is displaced by a water molecule in the high-humidity dataset (298 K, 99.5% RH), suggesting that the solvation distribution of the M^pro^ active site is malleable. Similarly, another DMSO in a distal region of the protein is ordered throughout the multitemperature series, but two waters and a new side-chain rotamer for Arg298 displace it in the high-humidity dataset.

Another putative active-site water, which we refer to as the “intervening water” or H_2_O_int_, is also present in each of our structures. However, unlike H_2_O_cat_, H_2_O_int_ is modeled in only <1% (2/252) of available structures: 7K3T (the highest-resolution M^pro^ structure available; apo state; B.A., D.K., M.R.F., S.M. *et al*., *in preparation*) and 7JFQ (“de-oxidized C145”, no publication). There are experimental differences among the structures that do have H_2_O_int_ modeled, as well as amongst those that do not have it modelled but do have electron density for it. For example, 7JFQ is in a different space group than our structures, with different crystallization conditions, including pH — so it is not immediately obvious what causes this water to sometimes be visibly ordered. See **Supp. Text 1** for a more thorough discussion of H_2_O_int_ modeling.

Interestingly, in our temperature series, H_2_O_int_ is not static, but rather varies in position across temperatures by nearly 1 Å (**Fig. 3a**), which is significantly in excess of the estimated coordinate error of 0.18–0.34 Å for our structures (calculated using the Diffraction Precision Index online server (Kumar *et al*., 2015)). Together with its absence in many other structures, this suggests that H_2_O_int_ is in some sense mobile. There is a rough trend of higher temperatures corresponding to H_2_O_int_ positions farther from the backbone of His41 and Cys145 and closer to His164, although this is not a strict rule. At 298 K, H_2_O_int_ is in an almost identical position regardless of humidity (ambient or 99.5% RH), suggesting both that environmental humidity has minimal effects on these crystals and again that our structural results with respect to temperature dependence may be precise. Some positional uncertainty may stem from the fact that the 2F_o_-F_c_ electron density for this water is not fully discrete, but rather appears semi-contiguous with the density for the adjacent catalytic His41 and Cys145 side chains (at typical map σ levels; **Fig. 2**).

**Figure 3:**
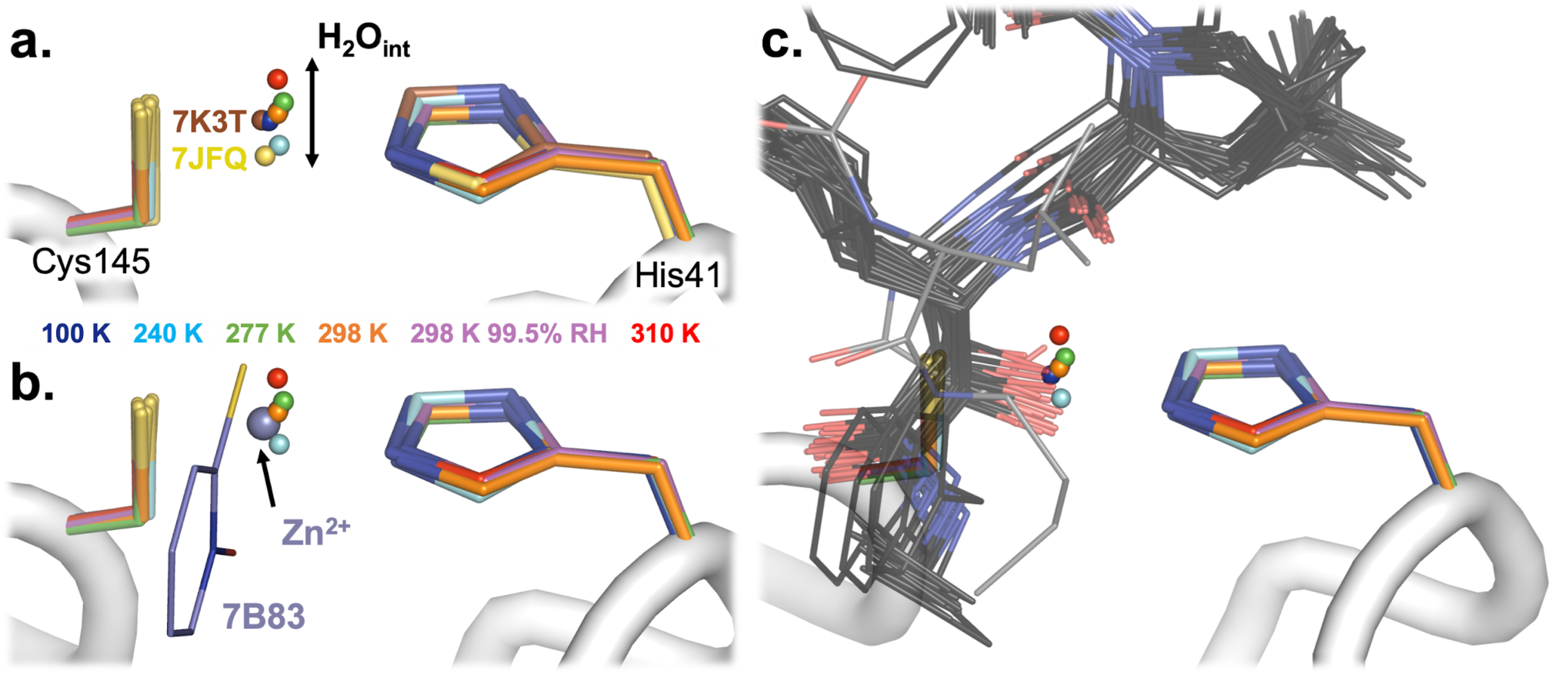
The rare active-site intervening water (H_2_O_int_) is sensitive to temperature and mimicked by ligands. **a**. In our new structures, H_2_O_int_ is ordered between Cys145 and His41 of the M^pro^ catalytic dyad, but its position varies as a function of temperature (blue to red). 298* K = 298 K at 99.5% relative humidity. The position of H_2_O_int_ in the 298 K model collected at 99.5% relative humidity occupies an extremely similar position as that of H_2_O_int_ in the 298 K model (orange). In only 2/252 previous structures of SARS-CoV-2 M^pro^ (7K3T and 7JFQ, both 100 K) is H_2_O_int_ also ordered. **b-c**. H_2_O_int_ is mimicked/displaced by particular atoms in other previous structures of M^pro^. Together these encompass the swath of H_2_O_int_ positions. **b**. In 7B83, Zn^2+^ from zinc pyrithione (purple) displaces H_2_O_int_. **c**. In several structures from different series of covalent ligands (grey) linked to Cys145, a hydroxyl oxygen of the covalent adduct displaces H_2_O_int_. One of these thiohemiketals is observed in a distinct *(R)* conformation (6XFN, lighter grey), which places the hydroxyl oxygen at a more extreme position corresponding to H_2_O_int_ in our 310 K structure. Together these binders approximate the swath of H_2_O_int_ positions in our multitemperature series.

Although H_2_O_int_ is absent from the vast majority of the other hundreds of crystal structures of M^pro^, it is displaced or “mimicked” by ligands in some instances. For example, a Zn^2+^ ion from an ionophore binds in the same position in PDB ID 7B83 (Günther *et al*., 2021) (**Fig. 3b**). In addition, Zn^2+^ alone was bound in two other structures, including PDB ID 7DK1 (Panchariya *et al*., 2021). However, Zn^2+^ is not consistent with our data (see **Supp. Text 1**). In addition, several covalent ligands that target the catalytic Cys145 form thiohemiketal or thiohemiacetal adducts, where the hydroxyl group is placed near His41 in a similar position as H_2_O_int_ in our structures (Sacco *et al*., 2020). (**Fig. 3c**). Interestingly, the swath of positions for this hydroxyl group, including a more extreme position due to a distinct conformation of the linker (PDB ID 6XFN), aligns with the swath of positions taken by H_2_O_int_ across our temperature series (plus PDB ID 7K3T and 7JFQ) (**Fig. 3c**). Together, these data suggest a structural niche for H_2_O_int_ that, regardless of its potential role in the catalytic mechanism, may be productively exploited for small-molecule ligand design.

Beyond the active site, we turned our attention to the nearby P5 binding pocket, specifically the loop composed of residues 192–198. Previously, the first report of a room-temperature structure of M^pro^, which was in the apo form (PDB ID 6WQF) (Kneller, Phillips, O’Neill, Jedrzejczak *et al*., 2020), noted that this loop adopted a different conformation than in a prior 100 K apo structure (PDB ID 6Y2E) (Zhang *et al*., 2020), including rotated peptide orientations for Ala194–Gly195 and Asp197–Thr198. However, all of the structures in our multitemperature series, including at lower temperatures (100 K and 240 K), have a single backbone conformation in this region that matches that of 6WQF (**Fig. 4a**). In addition, other apo cryogenic structures, including one at high (1.2 Å) resolution (PDB ID 7K3T), also match the 6WQF backbone conformation. All of these structures (6WQF, 6Y2E, 7K3T, and our multitemperature series) derive from the same crystal form (**Table 1**). Thus, it appears the different loop conformation adopted in 6Y2E is not driven by temperature (Kneller, Phillips, O’Neill, Jedrzejczak *et al*., 2020), nor by ligand binding or crystal lattice effects, but rather by some other aspect of the crystallization details or sample handling conditions — including, perhaps, idiosyncratic effects of crystal cryocooling (Halle, 2004; Keedy *et al*., 2014). Our conclusion here is also supported by a recent retrospective analysis of existing structures (Jaskolski *et al*., 2021).

**Figure 4:**
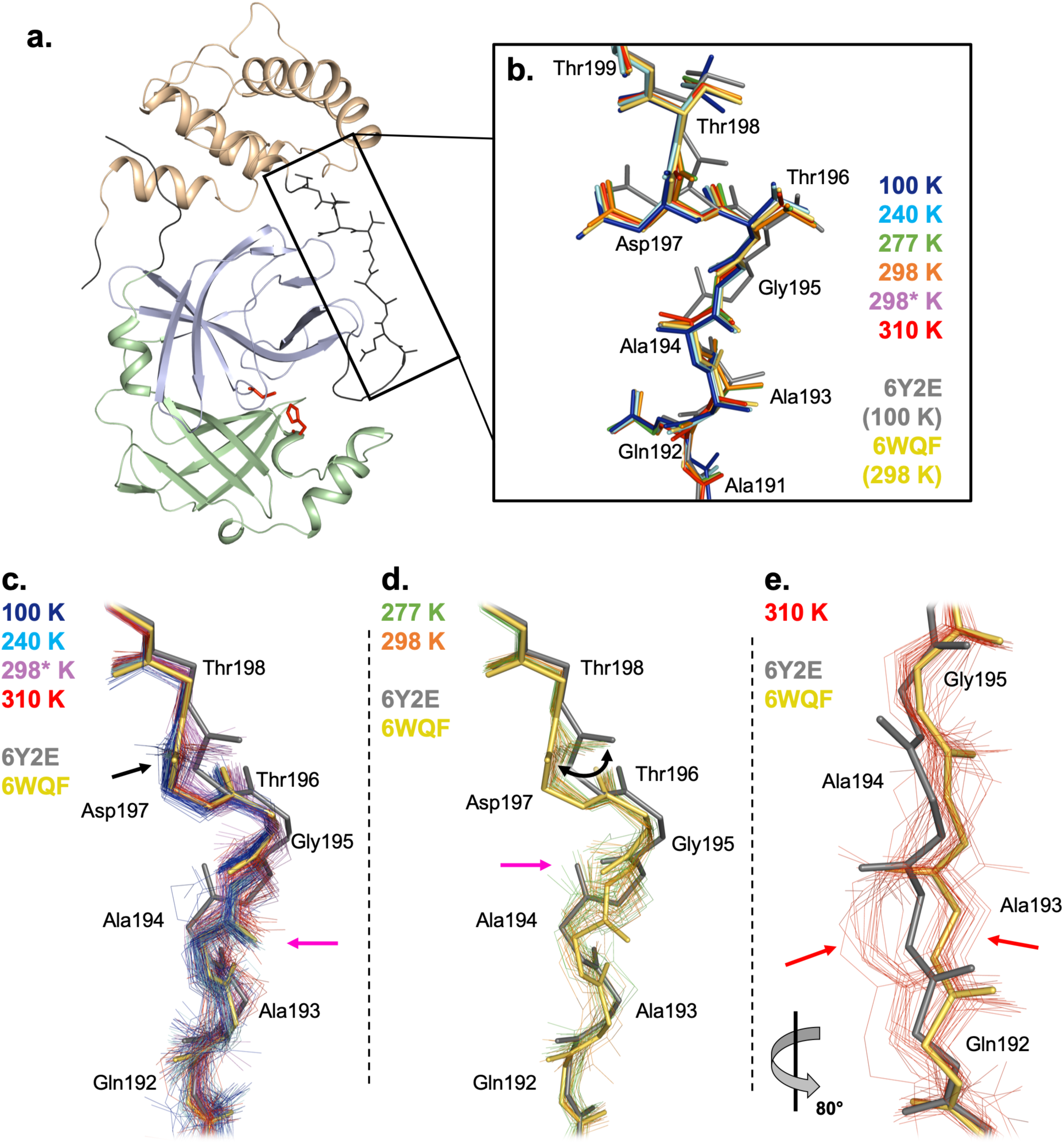
Complex temperature dependence of residues 192–198 in the P5 binding pocket. **a**. M^pro^ monomer from cryogenic structure, coloured by domain: domain I, residues 8–101, pale green; domain II, residues 102–184, pale blue; domain III, residues 201–303, pale orange. Catalytic dyad residues Cys145 and His41 are shown as sticks (red). Terminal residues are shown in dark grey. P5 binding pocket linker loop (residues 190–200) shown in dark grey and as sticks (black box). **b**. Our new multitemperature structures all have a single backbone conformation for this linker loop region. Regardless of temperature, they all match a similar backbone conformation to the room-temperature 6WQF model (yellow), and not the cryogenic 6Y2E model (grey) (Kneller, Phillips, O’Neill, Jedrzejczak *et al*., 2020). (298* K = 298 K, 99.5% relative humidity.) **c-e**. Phenix ensemble refinement models based on our multitemperature datasets reveal a complex pattern of flexibility that was “hidden” in **b**. **c**. For some conditions (100 K, blue; 240 K, cyan; 310 K, red; 298* K, magenta), the ensemble models generally match 6WQF, albeit with variability around the average conformation. For the Ala194–Gly195 peptide (pink arrow), all four conditions match 6WQF. For the Asp197–Thr198 peptide (black arrow), 100 K, 240 K, and 310 K match 6WQF, whereas 298* K adopts a swath of orientations at the Asp197–Thr198 peptide, bridging 6WQF and 6Y2E. **d**. For other conditions (277 K, green; 298 K, orange), the ensemble models exhibit shifts away from 6WQF and toward 6Y2E. For the Ala194–Gly195 peptide, both conditions match 6Y2E (pink arrow) instead of 6WQF. For the Asp197–Thr198 peptide, both conditions adopt a swath of orientations (black curved arrow) bridging 6WQF and 6Y2E, similarly to 298* K (in **c**). **e**. A zoomed-in and ∼80° rotated view of the 310 K ensemble model illustrates a split between the primary conformation and a distinct alternate conformation, centered on Ala193 (red arrows). This split is unique to the 310 K ensemble model.

### Crystallographic ensemble models reveal distinct backbone conformational heterogeneity

We next aimed to complement this analysis of our manually built multiconformer models with a more automated and explicitly unbiased approach to modeling flexibility that can handle larger-scale backbone flexibility such as loop motions. Therefore, we turned to Phenix ensemble refinement, which uses molecular dynamics simulations with time-averaged restraints to crystallographic data (Burnley *et al*., 2012). Phenix ensemble models have been used fruitfully for many applications (Woldeyes *et al*., 2014), including exploring the effects of temperature on protein crystals (Keedy *et al*., 2014), assessing the conformational plasticity of peptide–MHC interactions (Fodor *et al*., 2018), and rational protein design (Broom *et al*., 2020). After a scan of parameter space (see Methods), we created one ensemble model per temperature, each of which contains 18 to 45 constituent models (**Table 2**). Compared to the multiconformer models, the ensemble models fit the experimental data equally well or better based on R_free_, albeit with slightly wider R_free_-R_work_ gaps (**Table 2** vs. **Table 1**).

**Table 2:** Refinement statistics for Phenix ensemble models. p_TLS_ and w_x-ray_ are input parameters to Phenix ensemble refinement; the other input parameter (τ_x_) was automatically determined (see Methods).

| Structure | 100 K | 240 K | 277 K | 298 K | 298 K,<br>99.5% RH | 310 K |
| --- | --- | --- | --- | --- | --- | --- |
| PDB ID | 7MHL | 7MHM | 7MHN | 7MHO | 7MHP | 7MHQ |
| Resolution (Å) | 1.55 | 1.53 | 2.19 | 1.88 | 2.00 | 1.96 |
| $p_{\text{TLS}}$ | 1 | 1 | 1 | 1 | 1 | 1 |
| $w_{\text{x-ray}}$ | 2.5 | 2.5 | 2.5 | 2.5 | 2.5 | 2.5 |
| # models in ensemble | 45 | 43 | 18 | 20 | 36 | 42 |
| $R_{\text{work}}$ | 0.1700 | 0.1606 | 0.1626 | 0.1667 | 0.1592 | 0.1686 |
| $R_{\text{free}}$ | 0.2235 | 0.1978 | 0.2169 | 0.2077 | 0.2174 | 0.2240 |

Using these ensemble models, we reexamined the P5 binding pocket loop mentioned above. The 100, 240, and 310 K ensemble models are similar to the previous “room-temperature” structure 6WQF, with mostly the same peptide orientation for Ala194–Gly195 and Asp197–Thr198 (**Fig. 4c**). By contrast, the 277 and 298 K ensemble models match the flipped Ala194–Gly195 peptide orientation from the previous cryogenic structure 6Y2E, and sample a swath of conformations for Asp197–Thr198 that span 6WQF and 6Y2E (**Fig. 4d**). Interestingly, although the 298 K 99.5% RH model follows a similar Ala194–Gly195 peptide orientation to 100, 240 and 310 K, it also exhibits a swath of peptide conformations for Asp197–Thr198, similar to our 277 and 298 K ensemble models. The distinction between ensembles that match 6WQF vs. 6Y2E is not simply a byproduct of resolution: although 277 K has the lowest resolution (2.19 Å), 298 K (1.88 Å) has a better resolution than 310 K (1.96 Å).

Further, our 310 K ensemble reveals a distinct conformational split in this loop region, centered on Ala193, indicated by a subset of models within the ensemble following the primary conformation and a second subset following a separate, different conformation (**Fig. 4e**). Taken together, these results suggest that this region may have complex temperature dependence, as well as the capacity to sample alternate conformations that are “captured” in particular individual structures.

Beyond just the P5 loop, we also examined other regions with elevated and/or temperature-dependent ensemble Cα root-mean-square fluctuation (RMSF) (**Fig. 5a**) that were not previously noted as being temperature-dependent. These regions segregate into different categories with distinct temperature dependence.

**Figure 5:**
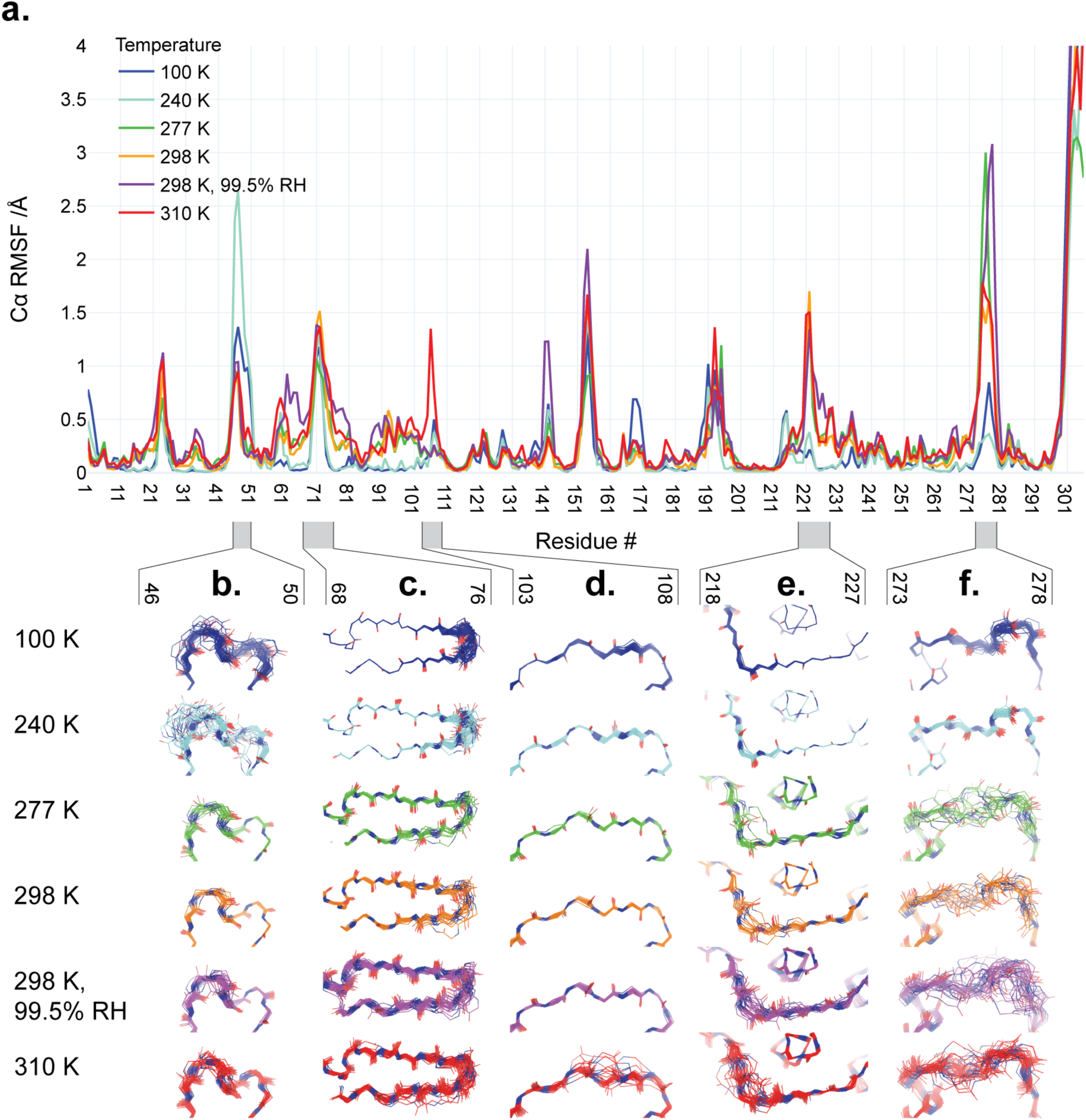
Backbone structural variability of ensemble models along the M^pro^ sequence as a function of temperature. **a**. Root-mean-square fluctuation (RMSF) of Cα backbone atom positions is plotted vs. residue number for each of the different structures in our multitemperature series (colors in legend). RMSF spikes at the N-terminus, C-terminus, and β-turn 153–157 (in contact with the C-terminus) in the ensemble models are truncated in this plot, and should be interpreted with caution. **b-f**. Backbone structures from ensemble refinement are shown for regions coinciding with temperature-dependent RMSF peaks. The refined single structure is shown as a cartoon, while atoms in the backbone of ensemble models are shown as lines.

First, some regions display a generally positive correlation between backbone structural variability and diffraction experiment temperature. For example, residues 218–227 (**Fig. 5e**) and 273–278 (**Fig. 5f**) are highly ordered at 100 K and 240 K, but mobile at warmer temperatures. These regions are spatially contiguous in the monomer, within the helical domain III. In another case, residues 68–76 (**Fig. 5c**), conformational diversity is restricted to the β-hairpin at 100 K and 240 K, but appears to spread further down the β-strands at higher temperatures. Interestingly, although 68–76 is isolated from the regions described earlier (218–227 & 273–278) in the monomer and the biologically-relevant dimer, it is contiguous with them in the crystal lattice. In contrast to these regions with quasi-continuous temperature dependence, we observe a more abrupt response for residues 103–108 (**Fig. 5d**), which shows significant backbone heterogeneity only at 310 K. Notably, this region is spatially separated (within the monomer, dimer, and lattice) from the regions mentioned above that have a less abrupt temperature response. Finally, we observe one region with an atypical relation between backbone variability and temperature: the short 3_10_ helix at residues 46–51 (**Fig. 5b**). This region abuts the P5 substrate binding loop composed of residues 192–198 with its complex temperature dependence (**Fig. 4**); together, these two regions form one side of the active site pocket (**Fig. 1b**).

### A network of coupled conformational heterogeneity bridges the active site, substrate pocket, inter-domain interface, and dimer interface

To complement the model-centric approaches above, we also looked for temperature-dependent conformational effects using an approach that is more directly data-driven: isomorphous F_o_-F_o_ difference electron density maps. We computed F_o_-F_o_ difference maps for each temperature vs. 100 K, and looked for patterns in terms of spatial colocalization of difference peaks. The global results confirm that the protein structure remains similar overall, with a smattering of difference peaks throughout the monomer asymmetric unit (**Supp. Fig. 4**). However, within those difference peaks lies a provocative stretch of difference features spanning the dimer interface, the interface between domain I and domain II of the monomer, and the edge of the P5 substrate binding pocket (**Fig. 6**). These difference features may be somewhat resolution-dependent, as they are least pronounced for 277 K (2.19 Å) and most pronounced for 240 K (1.53 Å), but their distribution across M^pro^ is qualitatively similar across temperatures.

**Figure 6:**
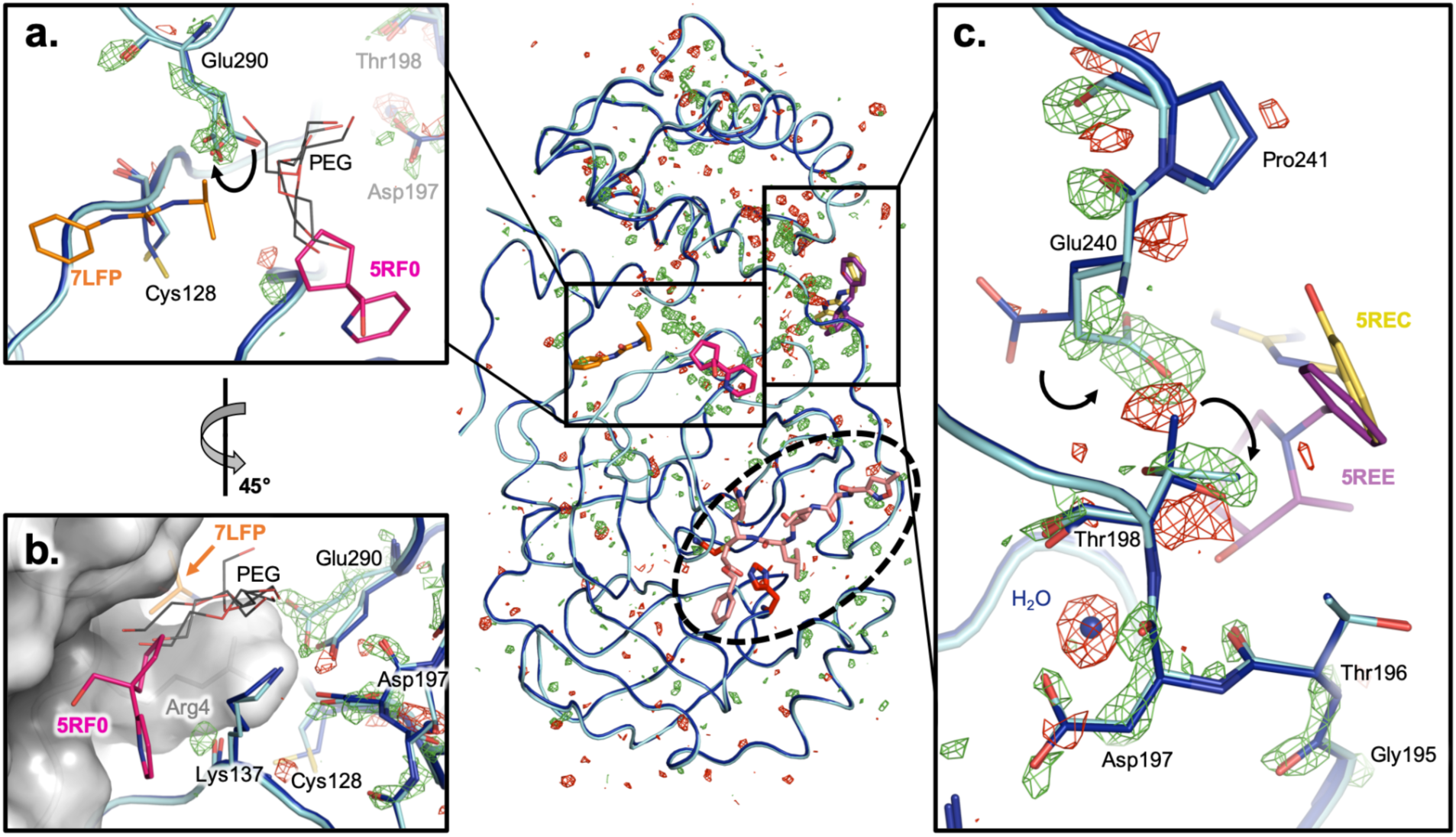
F_o_-F_o_ difference maps reveal local conformational shifts connecting the active site, inter-domain interface, and dimer interface. *Center:* Overview of isomorphous F_o_-F_o_ difference electron density map at +/-3 σ (green/red mesh) for the 240 K dataset (cyan) minus the 100 K dataset (dark blue). (See **Supp. Fig. 4** for F_o_-F_o_ maps for all temperatures). Ligands from cocrystal structures are shown at the active site (dashed oval) (pale orange, 6LU7), inter-domain interface (purple, 5REE; yellow, 5REC), and dimer interface (orange, 7LFP; pink, 5FR0). **a**. Glu290 switches from one side-chain rotamer at 100 K to two alternate rotamers at 240 K (curved arrow). Glu290 is spatially adjacent to Cys128, which switches from two alternate rotamers at 100 K to a single rotamer at 240 K in our multiconformer models. These residues are near two ligands from separate crystallographic screens (7LFP, 5RF0), as well as many ordered PEG molecules from the crystallization cocktails of various structures (7KVR, 7KVL, 7KFI, 7LFE). **b**. A ∼45° rotated view relative to **a**. shows that these two ligands bind at the dimer interface of the biological monomer, constituted in the crystal from a symmetry-related protomer (grey surface). This interface also includes the Asp197 region (right). **c**. Thr198 switches from two alternate side-chain rotamers at 100 K to a single rotamer at 240 K, while Glu240 — located across the inter-domain interface — changes side-chain rotamer (curved arrows), with additional effects on the adjacent backbone of Pro241. In the other direction from Asp197 (down in this view), other residues in the P5 substrate binding pocket loop (**Fig. 4**) undergo conformational adjustments en route to the active site. Meanwhile, an interacting water molecule at 100 K (blue sphere) becomes less ordered or displaced at 240 K.

A closer examination of the models in the vicinity of these difference features reveals what appears to be a series of correlated conformational motions keyed to temperature change. For example, F_o_-F_o_ density shows that Glu290 shifts from a single side-chain rotamer at 100 K to two alternate rotamers with partial occupancies at 240 K (**Fig. 6a**); this second rotamer seen at 240 K then remains as a single full-occupancy conformation for all higher temperatures. In sync with Glu290, our multiconformer models show that the adjacent Cys128 shifts its conformational distribution, but in the opposite fashion: from two alternate rotamers to one (**Fig. 6a**). Both Glu290 and Cys128 interact with a symmetry-related Arg4 across the biological dimer interface (**Fig. 6b**). Interestingly, two small-molecule fragments from recent crystallographic screens (Douangamath *et al*., 2020; Noske *et al*., 2021) bind at this area of the dimer interface (**Fig. 6a-b**). Moreover, ordered polyethylene glycol (PEG) molecules from several previous structures illustrate the potential for future ligand design efforts to “grow” from one of these initial fragment hits (5RF0) toward the mobile Glu290 and Cys128. This observation reinforces the idea that molecules from crystallization solutions, such as glycols, can reveal useful features like cryptic binding pockets (Bansia *et al*., 2021).

Glu290 is connected to another interesting residue, Asp197, via a hydrogen-bond network with only one intervening side chain (Arg131). Within this vicinity, an interacting water molecule is liberated, and an adjacent residue, Thr198, shifts from two alternate side-chain rotamers to just one (**Fig. 6c**). The Thr198 motion is linked to a conformational change for the nearby Glu240 side chain and Pro241 backbone, thus establishing a possible means for allosteric communication across the inter-domain interface. In the opposite direction from Asp197, other adjacent residues experience changes in ordering per F_o_-F_o_ peaks; these residues together form the 192–198 loop of the functionally important and mobile P5 pocket (**Fig. 4**) leading toward the active site.

Overall, these observations describe a series of conformational motions that bridge the dimer interface, inter-domain interface, substrate binding pocket, and active site (**Fig. 6** *center*, boxes and oval). In this work, temperature is the perturbation/effector — but our results raise the enticing possibility that future small molecules could be used to allosterically perturb this network, thereby modulating enzyme dimerization and/or catalysis.

## Discussion

Our crystal structures of unliganded SARS-CoV-2 M^pro^ at variable temperature and humidity paint a picture of a complex protein conformational landscape. The structure of M^pro^ does not change linearly with temperature; rather, there is a global transition between roughly <240 K and >277 K (**Fig. 1d, Supp. Fig. 2**). This 240–277 K transition regime for M^pro^ does not coincide with the 180–220 K glass transition or dynamical transition threshold seen previously for other systems such as CypA (Keedy *et al*., 2015), suggesting protein-to-protein variability. More locally in M^pro^, as temperature increases, different regions experience distinct types of changes to conformational heterogeneity (**Fig. 5**), in line with previous multitemperature studies of other proteins (Keedy *et al*., 2014). These effects are not limited to surface-exposed side chains as one might naïvely expect, but rather encompass motions of buried side chains (**Fig. 6**), many backbone regions (**Fig. 4, Fig. 5**), and water molecules themselves (**Fig. 2, Fig. 3**). Our results here for M^pro^, as well as a large body of previous literature for other systems, refute the assertion that X-ray crystallography under “unusual experimental conditions” like variable temperature is not useful for understanding proteins (Jaskolski *et al*., 2021). By contrast, our work is in line with computational analyses of B-factors suggesting that different alternate conformations for M^pro^ (and other systems) can be accessed by varying temperatures and/or the crystal lattice (Pearce & Gros, 2021).

A key example of temperature-sensitive, protein-associated solvent is the mobile water H_2_O_int_ that we observe in the M^pro^ active site (**Fig. 2, Fig. 3**) (see **Supp. Text 1**). This rarely observed water’s intriguing placement, with multiple permitted positions along a swath between the catalytic dyad of His41 and Cys145, suggests it may play some role in the catalytic process (Lee *et al*., 2020). Notably, recent structures of an acyl-enzyme intermediate structure (PDB ID 7KHP) and a C145A mutant product-bound structure (7JOY) of M^pro^ include a nearby water, ∼1.5 Å away but aligned with our approximately collinear multitemperature H_2_O_int_ swath (**Fig. 3**), which the authors suggested may play a role as a deacylating nucleophile (Lee *et al*., 2020). Questions about the functional role of H_2_O_int_ could be explored in parallel with other experiments to probe details of the catalytic mechanism, such as variable pH to probe Cys145 oxidation and reactivity (Kneller, Phillips, O’Neill, Tan *et al*., 2020) and neutron crystallography to reveal a zwitterionic state of the catalytic dyad (Kneller, Phillips, Weiss *et al*., 2020), although questions remain about the interpretation of such data (Jaskolski *et al*., 2021).

Perhaps surprisingly, unlike temperature, high relative humidity during data collection does not affect H_2_O_int_ in our structures (**Fig. 2, Fig. 3**). However, humidity does alter the solvation shell elsewhere nearby in the active site (**Fig. 2**, bottom right). Displaceable waters could potentially be exploited to design high-affinity small-molecule inhibitors, particularly when guided by water thermodynamics maps from simulations, as are available for M^pro^ and other SARS-CoV-2 proteins (Olson *et al*., 2020). More broadly, this result hints at the utility of humidity as an experimental variable in crystallography (Kiefersauer *et al*., 2000; Sanchez-Weatherby *et al*., 2009) for exploring solvent slaving to solvent energetics in ligand binding (Darby *et al*., 2019), protein dynamics (Lewandowski *et al*., 2015), and other functionally relevant phenomena.

Phenix ensemble models (Burnley *et al*., 2012) refined from our X-ray datasets helped us to illuminate temperature-dependent differences in conformational heterogeneity in certain areas of M^pro^ (**Fig. 4, Fig. 5**) that were concealed by more traditional model types (Babcock *et al*., 2018). Despite its utility in this and other work, there is significant potential for improvement of the ensemble refinement methodology through, for example, integration of more sophisticated molecular mechanics force fields like Amber (Moriarty *et al*., 2020) into the molecular dynamics component (Burnley *et al*., 2012) to improve ensemble model geometry, or more sophisticated treatments of translation-libration-screw (TLS) groups to isolate interesting local conformational heterogeneity (Ploscariu *et al*., 2021). Although it was also beyond the scope of this study, ensemble models may reveal alternate conformational substates that are important for the catalytic cycle, which could be fruitfully targeted by small molecules for antiviral drug design.

Finally, our results emphasize the allure of allosteric inhibition of M^pro^ as an alternative therapeutic strategy. Our structures illustrate apparently coupled conformational motions that bridge the active site, substrate binding pocket, inter-domain interface, and parts of the broad dimer interface (**Fig. 5, Fig. 6**). This is particularly noteworthy since M^pro^ must dimerize to become an active enzyme (Fan *et al*., 2004; Goyal & Goyal, 2020); inter-domain flexing has also been observed, even in crystals (Jaskolski *et al*., 2021). The intramolecular network we describe includes several sites that are distal from the active site, one of which is highlighted by unambiguous Glu240 difference density (**Fig. 6c**) corresponding to a temperature-dependent rotamer flip—this site has already been characterized as ligandable by recent crystallographic screens of pre-existing drug molecules (Günther *et al*., 2021) and small-molecule fragments (Douangamath *et al*., 2020) (**Fig. 6**). Some new M^pro^ ligands have been shown by mass spectrometry to disrupt the M^pro^ dimer and allosterically inhibit catalysis, albeit weakly thus far (El-Baba *et al*., 2020), illustrating the potential of an allosteric strategy. As a complementary structure-based approach to current experiments on the dimeric crystal form of M^pro^, future experiments could exploit mutations of the dimer interface to stabilize an inactive monomer, thus capturing a new structural target for crystallographic and solution screening for allosteric inhibitors that block dimerization. Ultimately, the present study offers insights into fundamental aspects of protein structural biophysics, and may also help pave the way for new efforts toward allosteric modulation of M^pro^ as a strategy for COVID-19 antiviral drug design.

## Methods

### Cloning, expression, and purification

Full details of the cloning, expression, and purification will be reported elsewhere (B.A., D.K., M.R.F., S.M. *et al*., *in preparation*). Briefly, the codon-optimized synthetic gene of full-length M^pro^ from SARS-CoV-2 was cloned into the pET29b vector. The cloned M^pro^ with C-terminal 6x histidine tag was expressed in *E. coli* using an auto-induction procedure (Studier, 2005). Cells were harvested, lysed using bacterial protein extraction agents (B-PER, ThermoFisher Scientific) in the presence of lysozyme, and purified with nickel-affinity chromatography followed by size exclusion chromatography. The histidine tag was cleaved by human rhinovirus (HRV) 3C protease (AcroBIOSYSTEMS) and further purified by reverse nickel-affinity chromatography. The purified protein was then dialysed overnight at 4°C against 30 mM HEPES pH 7.4, 200 mM NaCl, 1 mM TCEP; concentrated to ∼7 mg/mL; and used for crystallization or stored at 80°C.

### Crystallization

Plate-like crystals ranging from ∼100–400 µm along the longest axis (∼5–10 µm along the shortest axis) were grown via sitting drop vapor diffusion. The crystals grew in flower-like clusters (**Supp. Fig. 5**). After mixing a 1:1 ratio of ∼7 mg/mL M^pro^ with a solution of 22% PEG 4000, 100 mM HEPES pH 7.0, 3–5% DMSO and incubating at a temperature of ∼298 K, crystals were seen after 2–6 days.

### Crystal harvesting and X-ray data collection

Individual crystals were harvested using 10 µm MicroMesh™ loops (MiTeGen). For cryogenic temperature, crystals were cryocooled by the traditional practice of plunging into liquid nitrogen. For non-cryogenic temperatures at ambient humidity, crystals were coated with paratone oil, then mounted on the goniometer for data collection. Datasets were also collected for crystals coated with paratone oil and additionally enclosed in MicroRT™ capillaries (MiTeGen), but no differences were observed relative to paratone oil only. For high humidity, crystals were not coated with paratone oil, but were enclosed in MicroRT™ capillaries for the short transit to the goniometer, then removed once humid air flow was established on the goniometer; this ensured the crystal was always maintained at high humidity after leaving the crystallization drop. Each crystal was equilibrated on the goniometer for 10-20 minutes, more than sufficient to reach stable conditions.

X-ray data were collected at the National Synchrotron Light Source II (NSLS-II) beamline 17-ID-2 (FMX) (Schneider *et al*., 2021) using an X-ray beam of energy 12.66 keV, corresponding to a wavelength of 0.9793 Å; a horizontal-bounce Si111 double crystal monochromator; and an Eiger X 16M pixel array detector (Dectris). Temperature at the sample goniometer was controlled using a Cryostream 800 (OxfordCryosystems). For the 298 K, 99.5% relative humidity dataset, RH was controlled with an HC-LAB Humidity Controller (Arinax). Ambient temperature was measured to be ∼298 K, and ambient humidity was measured to be 36.7%. A new crystal was used for each dataset. Helical/vector data collection was used to traverse the length of each crystal, with a beam size of 10 × 10 µm. Using RADDOSE-3D (Bury *et al*., 2018), we estimated diffraction-weighted dose (DWD) for our datasets to be 242 kGy for 100 K, 532 kGy for 240 K, 397 kGy for 277 K, 137 kGy for 298 K, 182 kGy for 298 K (99.5% RH), and 176 kGy for 310 K. All of these DWD values are at or below the estimated room-temperature limit of about 400 kGy (Fischer, 2021) for our higher temperatures, although this limit is generally system-dependent. The DWD for 240 K is above the room-temperature limit, but such lower temperatures have higher dose tolerance. Additionally, there was no evidence of global radiation damage from R_d_ plots (**Supp. Fig. 1**), and local/specific radiation damage did not appreciably accrue during the course of each single-crystal data collection, as indicated by 2F_o_-F_c_ electron density maps around carboxyl groups (not shown).

### X-ray data reduction and modeling

The data reduction pipeline fast_dp (Winter & McAuley, 2011) was initially used for bulk data reduction during the beamtime, with selected data reprocessed using the xia2 DIALS (Winter *et al*., 2018) and xia2 3dii (XDS and XSCALE) pipelines (Kabsch, 2010), with xia2 3dii (XDS and XSCALE) also used for the generation of R_d_ statistics (Diederichs, 2006) (**Supp. Fig. 1**). Molecular replacement for each dataset was performed via Phaser-MR from the Phenix software suite, using PDB ID 6YB7 as a search model. Phenix AutoBuild (Terwilliger *et al*., 2008) was used for initial model building and refinement, with subsequent iterative refinements performed using phenix.refine (Afonine *et al*., 2012) and Coot (Emsley & Cowtan, 2004). After a few initial rounds of refinements, hydrogens were added using phenix.ready_set (Reduce (Word *et al*., 1999) and eLBOW (Moriarty *et al*., 2009)). For refinement of each dataset, X-ray/stereochemistry weight and X-ray/ADP weight were refined and optimised. Geometric and protein statistics of the final models were evaluated via MolProbity (Chen *et al*., 2010; Williams *et al*., 2018) and the JCSG-QC check server (https://smb.slac.stanford.edu/jcsg/QC/). Data collection and refinement statistics are shown in **Table 1**.

Crystallographic ensemble models were generated using phenix.ensemble_refinement (Burnley *et al*., 2012) in version 1.18.2-3874 of Phenix. Alternate conformations were first removed from the multiconformer models, and hydrogens were (re)added using phenix.ready_set. Next, a phenix.ensemble_refinement grid search was performed by repeating the simulation with four values of p_TLS_ (1.0, 0.9, 0.8, 0.6) and three values of wxray_coupled_tbath_offset (10, 5, 2.5), and using a random_seed value of 2679941. τ_x_ was set automatically according to the high-resolution limit of the dataset. From this grid, we present the analysis of the set of ensemble models that has both the lowest mean R_free_ and the lowest mean R_free_-R_work_ gap: p_TLS_=1.0, wxray_coupled_tbath_offset=2.5. Conclusions drawn for the set of ensemble models with lowest R_free_ per dataset, or the lowest R_free_-R_work_ gap per dataset, were similar. Refinement statistics are shown in **Table 2**.

For F_o_-F_o_ isomorphous difference map analysis, the phenix.fobs_minus_fobs_map executable in the Phenix software suite was used. Each elevated temperature was compared to 100 K. The 100 K multiconformer model was used for phasing for each difference map. For solvent content analysis, rwcontents v7.1.009 from the CCP4 suite (Winn *et al*., 2011) was used.

## Accession numbers and data availability

Models and structure factors are available in the Protein Data Bank under the following PDB ID accession codes (see also **Table 1** and **Table 2**): **<u>7MHF</u>, <u>7MHG</u>, <u>7MHH</u>, <u>7MHI</u>, <u>7MHJ</u>, <u>7MHK</u>** for muticonformer models, and **<u>7MHL</u>, <u>7MHM</u>, <u>7MHN</u>, <u>7MHO</u>, <u>7MHP</u>, <u>7MHQ</u>** for ensemble models.

Diffraction data are available at the Integrated Resource for Reproducibility in Macromolecular Crystallography (https://proteindiffraction.org) under the following Digital Object Identifier names: **<u>10.18430/m37mhf</u>, <u>10.18430/m37mhg</u>, <u>10.18430/m37mhh</u>, <u>10.18430/m37mhi</u>, <u>10.18430/m37mhj</u>, <u>10.18430/m37mhk</u>**.

## Glossary

F_o_-F_o_: isomorphous difference electron density map
COVID-19: coronavirus disease 2019
SARS-CoV-2: severe acute respiratory syndrome coronavirus 2
M^pro^: SARS coronavirus main protease

