## Supplemental Figures and Text for "The temperature-dependent conformational ensemble of SARS-CoV-2 main protease (M^pro^)"

### Supplementary Information

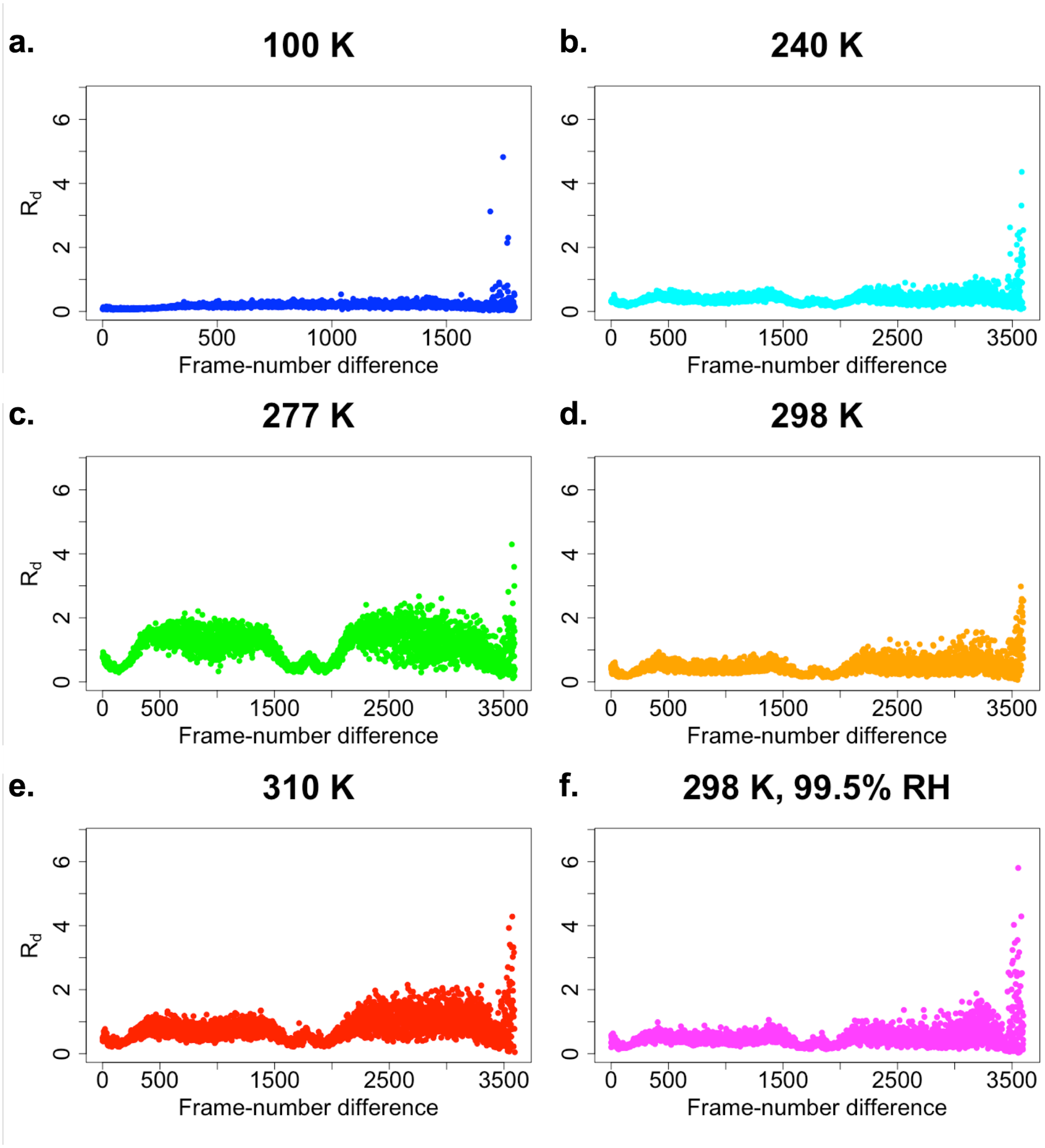


**Supplementary Figure 1:** Global radiation damage does not noticeably accrue during the course of each single-crystal data collection, as indicated by $R_d$ as a function of frame-number difference. Symmetric shapes of the plots derive from rotation of the thin, plate-like crystals during data collection, which unavoidably leads to different crystal volumes being irradiated for different frames.

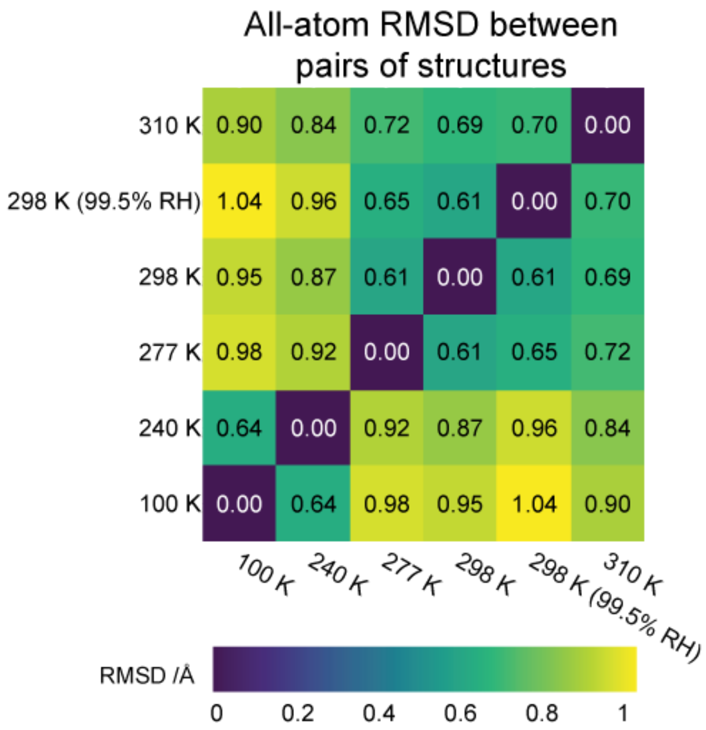


**Supplementary Figure 2:** Heatmap of pairwise all-atom RMSD between final refined structures, revealing temperature-dependent clustering (top-right vs. bottom-left).

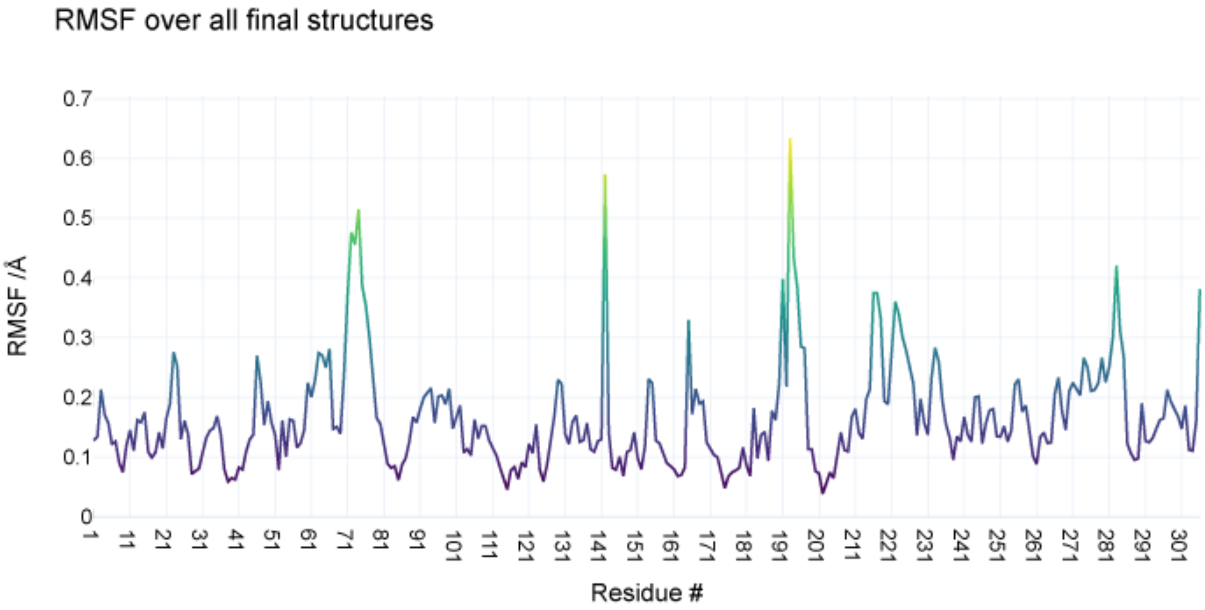


**Supplementary Figure 3:** Root-mean-square fluctuations (RMSF) of Cα atom positions over final refined structures. See also **Fig. 1c**.

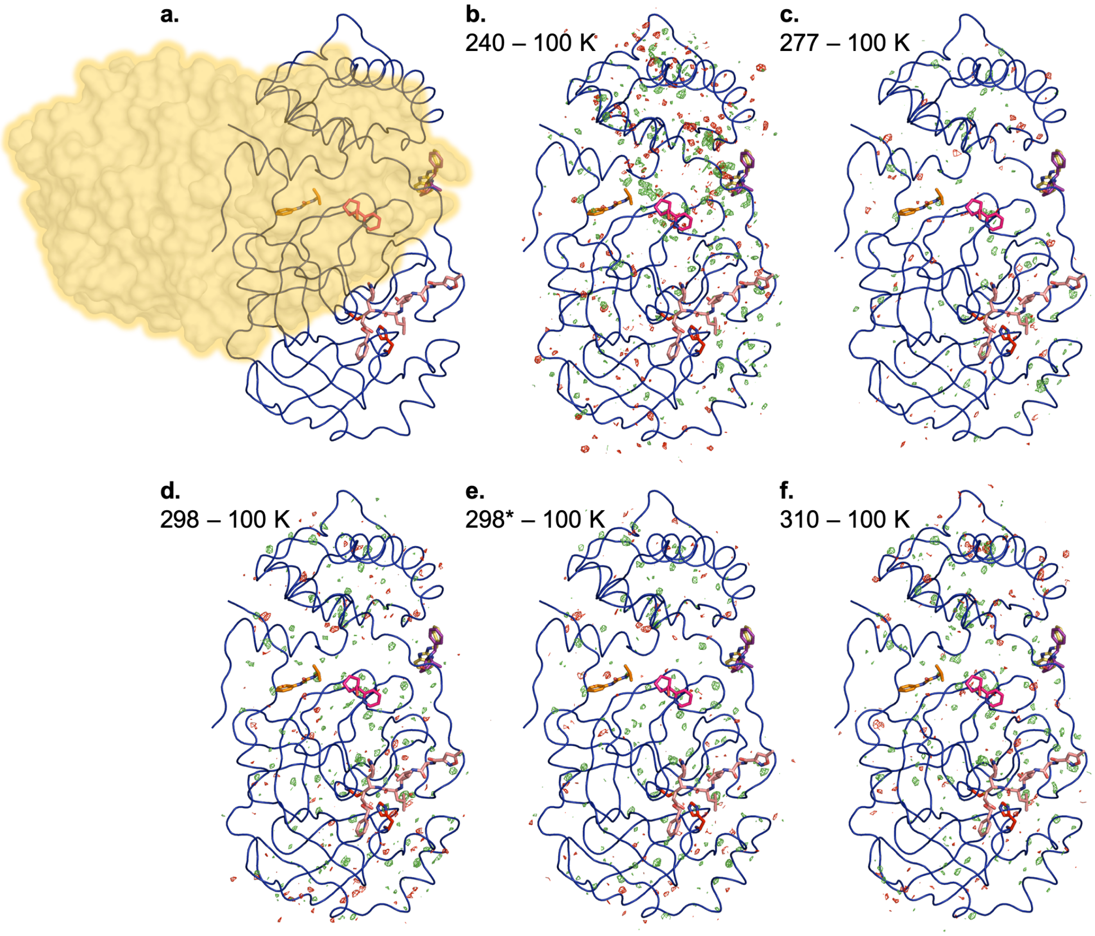


**Supplementary Figure 4:** Isomorphous $F_o$-$F_o$ difference electron density maps for all elevated temperatures. Ligands from cocrystal structures are shown as sticks at the active site (pale orange, 6LU7), inter-domain interface (purple, 5REE; yellow, 5REC), and dimer interface (orange, 7LFP; pink, 5FR0). **a.** Overview of 100 K structure with superimposed ligands, and highlighted dimer interface (orange). **b-f.** $F_o$-$F_o$ difference maps (+/-3 σ, green/red) are shown for each temperature minus 100 K.

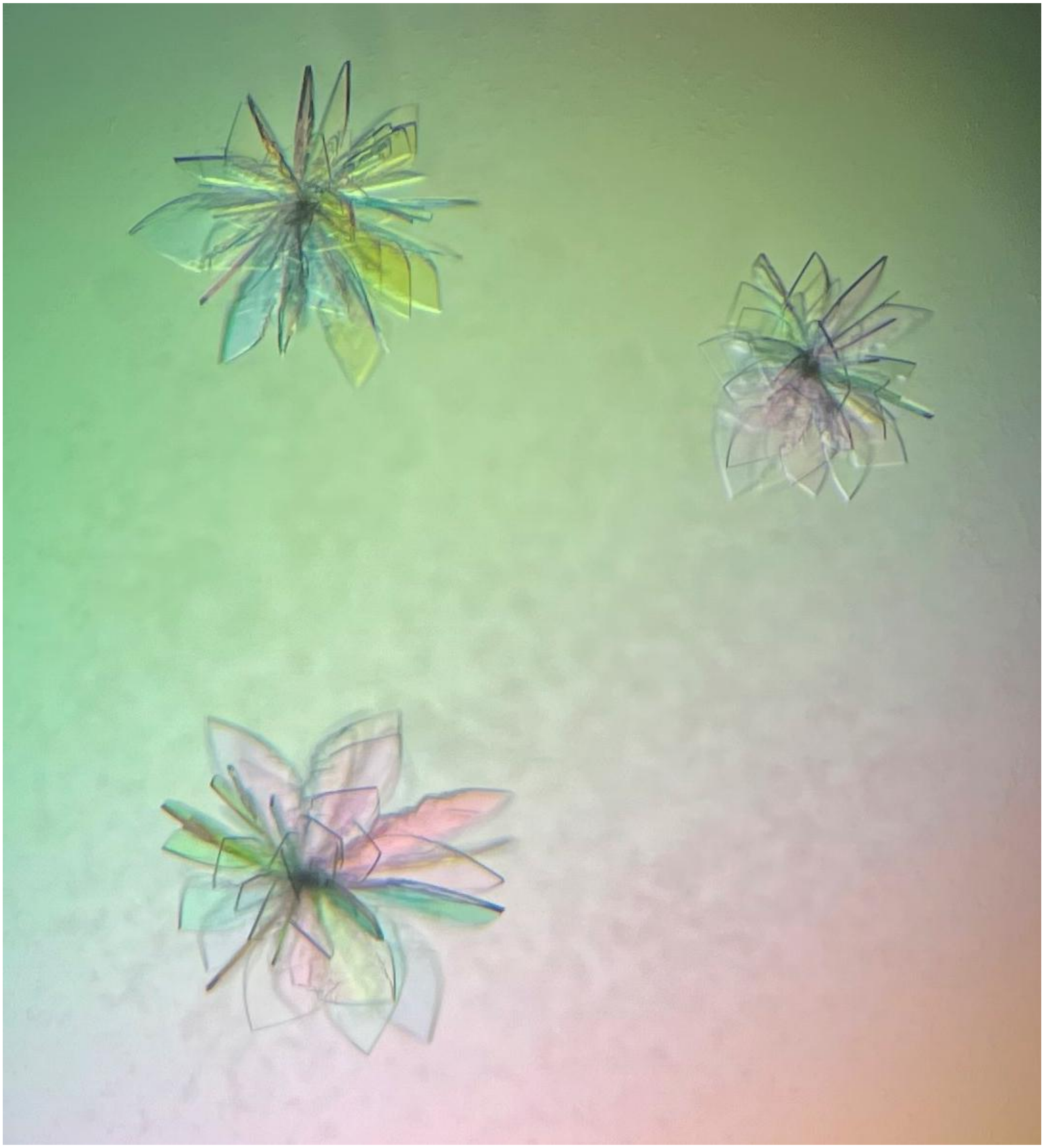

**Supplementary Figure 5:** Flower-like clusters of $M^{pro}$ crystals. For each dataset, one “flower petal” (crystal) was manually harvested and used for X-ray diffraction. Each petal is approximately ~100–400 µm along the longest axis and ~5–10 µm along the shortest axis.

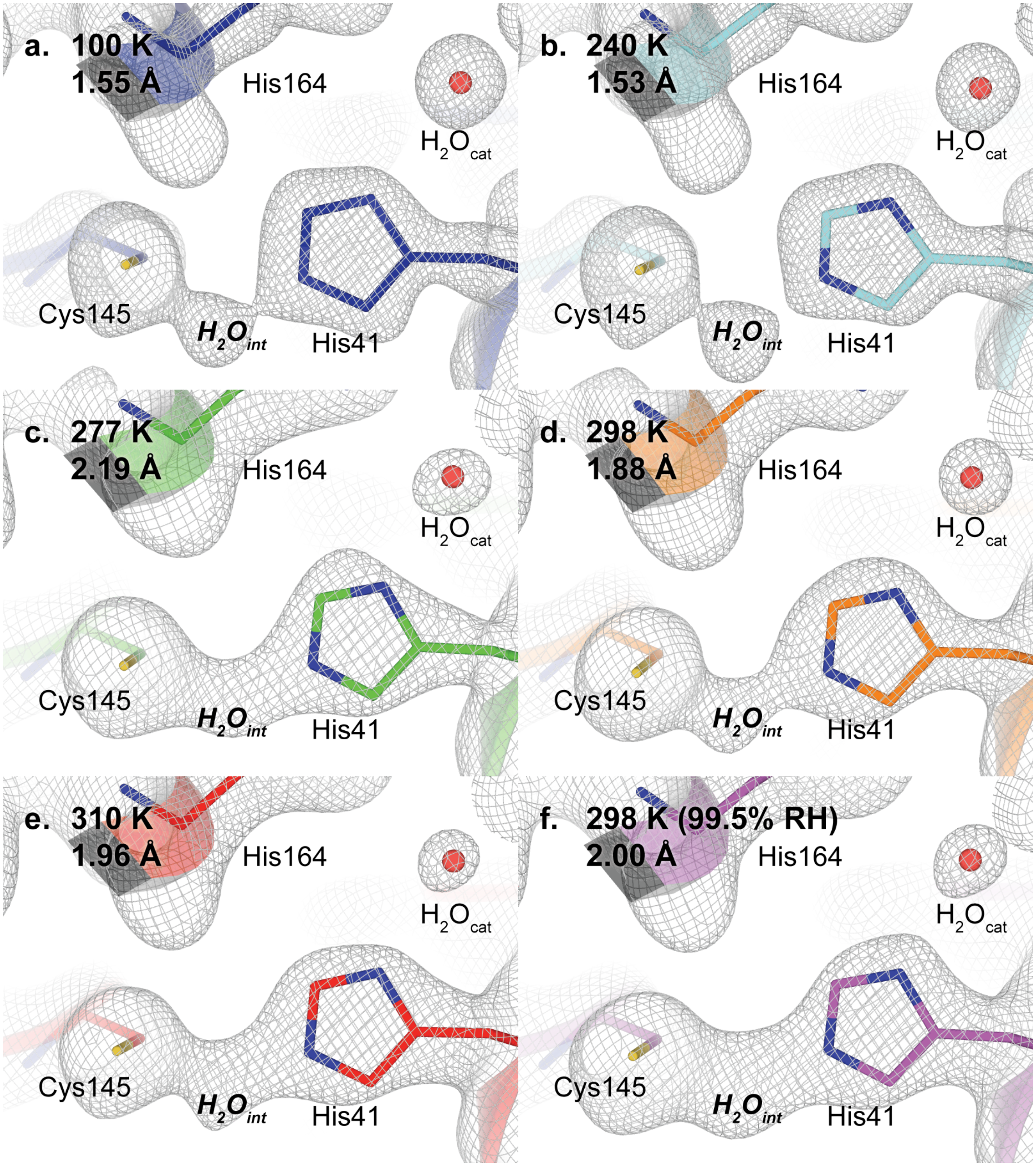


**Supplementary Figure 6:** Composite omit map centered on the active site, contoured at 1 σ. A region of density is observed between His41 and Cys145.

**Supplementary Text 1:** *Rationale for modeling $H_2O_{int}$.* The electron density maps from the X-ray datasets in this work reveal very clear evidence for an atom present in between Cys41 and His145 of the catalytic dyad, with relatively short distances to both (**Fig. 2**, **Supp. Fig. 6**). We have chosen to model this density as a water molecule, which we call $H_2O_{int}$ (**Fig. 3**). Here we discuss several relevant considerations, alternative modeling possibilities, and the reasoning behind our modeling choice.

<u>Presence of the density</u>

- Similar electron density is present and modelled in two other previously deposited structures, as discussed in the main text. One of these, PDB ID 7K3T, a cryogenic crystal structure of apo $M^{pro}$, is the highest-resolution structure of $M^{pro}$ yet determined: at 1.2 Å, it is at least 0.35 Å higher-resolution than any of our new structures. In 7K3T, the density is very clear in this region, and it is modeled as a water.
- For thoroughness, we also looked for unmodelled electron density in the PDB. We searched the PDB using a sequence similarity search at 97% identity to 7K3T chain A (381 structures as of 2021-Oct-10), filtered to X-ray diffraction, with structure factors, unrelated to PanDDA group depositions or ensembles (252 structures).
  By examining the published $2F_o$-$F_c$ maps of each of these 252 structures, we found density at the $H_2O_{int}$ site in 7 datasets other than ours (6M2Q, 6XB2, 6XKF, 7JFQ, 7JVZ, 7LMD, 7K3T).
  To reduce bias, a simple composite omit map was made for these datasets: there is clear density at the $H_2O_{int}$ site in three previously deposited apo datasets (6M2Q (unmodelled), 7JFQ, 7K3T), in addition to ours.
  These three datasets were collected across two different crystal packings (7JFQ: $P\ 2_1\ 2_1\ 2$, others C 2), and a range of crystallization pHs (range 5.8−8.5). Thus, we find it unlikely that this density is an artifact of conditions such as crystallization pH.

<u>Identity of the density</u>

- We used the Phenix automated ion/metal identification tool in phenix.refine (Echols *et al.*, 2014) to assess alternatives for the identity of $H_2O_{int}$ for all of our new datasets. The machine learning model from this tool did not flag $H_2O_{int}$ as a likely candidate for an ion/metal at any temperature or humidity.
- $Ca^{2+}$: For the 7K3T structure, a calcium complex was soaked into the crystal before data collection (B.A., D.K., M.R.F., S.M. *et al., in preparation*). However, the density for $H_2O_{int}$ looks very similar in 7K3T vs. our structures, which were not soaked with any calcium. PDB ID 6YVF has calcium modeled, but not in the active site. This suggests that calcium binds to $M^{pro}$, but elsewhere in its structure. For these reasons, calcium is an unlikely candidate for the $H_2O_{int}$ density.
- $Mg^{2+}$: Similarly, PDB ID 6WTT has magnesium modeled in $M^{pro}$, but not in the active site. Thus, magnesium is also an unlikely candidate.
- $Ni^{2+}$: It is formally possible that nickel ions from the affinity column bound to $M^{pro}$ during purification and remained bound throughout our experiments. However, in other structures that also used nickel affinity columns, density for $H_2O_{int}$ is not observed—either at cryogenic temperature (PDB ID 6Y2E, among others) or room temperature (PDB ID 6WQF, albeit at lower resolution). Therefore, exogenous nickel is an unlikely candidate for the $H_2O_{int}$ density.
- $Na^+$: Sodium was present in the final crystallization buffer, so it is a possibility for the identity of this density. However, we would expect a $Na^+$ to be bound by an oxygen atom on the

protein, and have either 5 or 6 neighbors (Harding, 2002). Furthermore, the Phenix automated ion picker did not score $Na^+$ strongly. While $Na^+$ and $H_2O$ are difficult to distinguish owing to their number of electrons, we do not believe that the lines of evidence available to us favor modelling $Na^+$ over $H_2O$.

- $Zn^{2+}$: Zinc is a possible candidate for the $H_2O_{int}$ density. Zinc often binds to Cys and His residues in proteins. Zinc binds tightly to $M^{pro}$, with 300 nM affinity (Panchariya *et al.*, 2021). Also, in previous structures of $M^{pro}$ with zinc bound (e.g. PDB ID 7DK1, 7D64) (Panchariya *et al.*, 2021), the zinc matches the position of our $H_2O_{int}$ density. However, those structures derive from crystals that were soaked in an excess of zinc, whereas we did not add any metals to our crystals (although it is formally possible that endogenous zinc from *E. coli* adhered to $M^{pro}$ during expression/purification). In addition, previous zinc-bound structures like 7DK1 have electron density for two additional waters to complete a tetrahedral coordination shell around the zinc (Panchariya *et al.*, 2021), whereas we do not see electron density in these positions even at low contour levels. We have also attempted modeling zinc in place of $H_2O_{int}$ in our structures, but refinement of such models results in strong negative $F_o$-$F_c$ difference density peaks, suggesting something as heavy as zinc is not present at this position. For these reasons, zinc remains a possibility in our structures, but several lines of evidence argue against it.
- Somewhat relatedly, although several nearby waters have weaker density and higher B-factors than $H_2O_{int}$, the nearby $H_2O_{cat}$ (**Fig. 2**) has stronger density and a lower B-factor than $H_2O_{int}$ and is indisputably a water. This suggests that $H_2O_{int}$ may indeed be a water as well, as opposed to a heavier metal.

Choice of water

- According to MolProbity, $H_2O_{int}$ in our models has steric clashes with Cys145 and His41. However, it is possible that Cys145 is deprotonated and anionic, and His41 is protonated at the Nε2 atom and cationic. Such a zwitterionic configuration was observed recently by neutron crystallography for apo $M^{pro}$ (Kneller, Phillips, Weiss *et al.*, 2020). In this scenario, $H_2O_{int}$ could be considered a hydroxide ion ($OH^-$) that is engaged in atypical, stronger, charge-assisted H-bonds (Gilli & Gilli, 2000). Although this interpretation is alluring, it is beyond the scope of our current variable-temperature/humidity X-ray crystallography study to perform additional neutron crystallography experiments to investigate protonation states in our crystals.
- We briefly considered leaving the $H_2O_{int}$ density unmodeled, but decided against it. In our view, leaving such strong density unmodeled would constitute an error of omission. Many crystal structures, both in general and particularly for SARS-CoV-2 $M^{pro}$, are used downstream by computational and medicinal chemists who do not inspect the original data or electron density maps; for these structure users, a model lacking any atom at the $H_2O_{int}$ position would be misleading.

Graphical Abstract

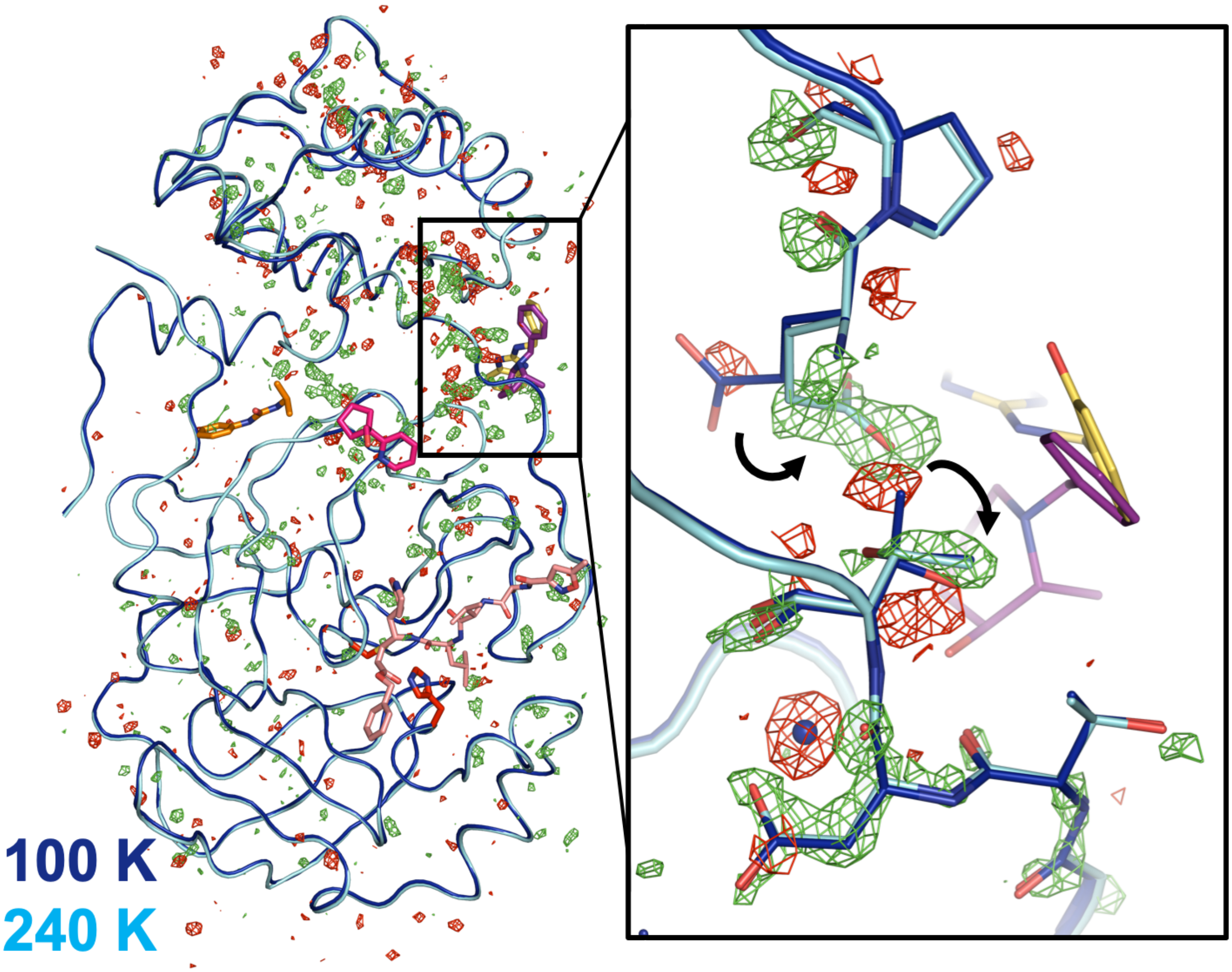
